# Interpreting *dN/dS* under different selective regimes in cancer evolution

**DOI:** 10.1101/2021.11.30.470556

**Authors:** Andrés Pérez-Figueroa, David Posada

## Abstract

The standard relationship between the *dN/dS* statistic and the selection coefficient is contingent upon the computation of the rate of fixation of non-synonymous and synonymous mutations among divergent lineages (substitutions). In cancer genomics, however, *dN/dS* is typically calculated by including mutations that are still segregating in the cell population. The interpretation of *dN/dS* within sexual populations has been shown to be problematic. Here we used a simple model of somatic evolution to study the relationship between *dN/dS* and the selection coefficient in the presence of deleterious, neutral, and beneficial mutations in cancer. We found that *dN/dS* can be used to distinguish cancer genes under positive or negative selection, but it is not always informative about the magnitude of the selection coefficient. In particular, under the asexual scenario simulated, *dN/dS* is insensitive to negative selection strength. Furthermore, the relationship between *dN/dS* and the positive selection coefficient depends on the mutation detection threshold, and, in particular scenarios, it can become non-linear. Our results warn about the necessary caution when interpreting the results drawn from *dN/dS* estimates in cancer.

## Introduction

The identification of genomic regions under natural selection plays a central role in evolutionary biology. As a consequence, a plethora of statistical tests have been proposed to identify and quantify evolutionary pressures at the molecular level. One of the most popular statistics is the *dN/dS* ratio, which compares the rate of synonymous substitutions (*dS*), assumed to be neutral, with the rate of non-synonymous substitutions (*dN*), which result in amino acid changes and can be targeted by selection. *dN/dS* is expected to be above one if selection promotes changes in the protein (positive selection), below one when it suppresses them (negative selection), and around one when protein changes are not favored or disfavored (neutrality). Importantly, this interpretation of *dN/dS* in terms of selective pressure is based on the comparison of substitutions (i.e., fixed differences) among distantly related lineages (Goldman and Yang, 1994; Li et al., 1985; Miyata and Yasunaga, 1980; Muse and Gaut, 1994; Nei and Gojobori, 1986), for which the relationship between *dN/dS* and the selection coefficient can be approximated as a deterministic function (Kryazhimskiy and Plotkin, 2008; Nielsen and Yang, 2003). Despite this requisite, *dN/dS* is often estimated at the population level, where changes among lineages do not represent only substitutions but also polymorphisms that are still segregating in the population. In this case, the dynamic of *dN/dS* is fairly distinct from the one assumed for interspecific comparisons (Fay, 2011; Kryazhimskiy and Plotkin, 2008; McDonald and Kreitman, 1991). Notably, within populations, the relationship between *dN/dS* and the selection coefficient is not always a monotonic function, and therefore it may be unfeasible to infer selective pressures from *dN/dS* (Kryazhimskiy and Plotkin, 2008). Furthermore, at the short time-scales typical of closely-related species and populations, the contribution of segregating polymorphisms induce a time-dependence factor in the estimation of *dN/dS*, particularly for single gene estimates, where the low number of mutations adds further noise (Mugal et al., 2020, 2014).

The *dN/dS* metric is widely used to identify genes under selection in cancer (Colom et al., 2020; Lawson et al., 2020, 2020; Martincorena et al., 2018, 2017; Tilk et al., 2019; Van den Eynden and Larsson, 2017; Wu et al., 2016; Zapata et al., 2020, 2018) and normal tissues (Colom et al., 2020; Lawson et al., 2020; Martincorena et al., 2018; Williams et al., 2020). Tumors are expanding masses of cells, and most of the mutations identified when sequencing a tumor sample are still segregating in the cell population (in the somatic/cancer jargon, these are *subclonal* mutations). Even those mutations that seem to be fixed (*clonal* mutations) may not be genuine substitutions because of the large sampling bias imposed by tissue biopsies (Chkhaidze et al., 2019). Furthermore, we should consider the nature of somatic evolution needs in the interpretation of *dN/dS*. For example, recombination is absent –or almost absent– potentially confounding positive and negative selection (Fay, 2011). Sampling time is another relevant factor to estimate the selective pressures in somatic tissues. It is well-known that in asexual and closely related populations, *dN/dS* changes over time because of a lag in the removal of slightly deleterious non-synonymous mutations (Mugal et al., 2020, 2014; Rocha et al., 2006). Recently, Williams et al. (2020) showed, using data from mutant clones in the normal esophageal epithelium (Martincorena et al. 2018) and normal skin (Martincorena et al. 2015), that the magnitude of the selection coefficient in somatic tissues is not necessarily well represented by a point estimate of *dN/dS* that ignores the frequency of the subclonal mutations.

To further assess the potential limitations of *dN/dS* for the study of cancer evolution, here we carried out computer simulations to *i*) describe the relationship of *dN/dS* with the selection coefficient during tumor growth, and *ii*) evaluate the accuracy by which linked neutral, deleterious and driver genomic regions can be identified from cancer patient cohorts using this statistic. Our results suggest that *dN/dS* can be used safely in cancer to differentiate genes under overall negative or positive selection or effectively neutral. On the other hand, the value of *dN/dS* depends to a large extent on the degree of polymorphism present at the time of sampling and the allele frequency threshold applied for its computation, resulting in some cases in a non-linear relationship between *dN/dS* and the selection coefficient.

## Methods

### Simulation model

We generated *in silico* tumor cohorts using OncoSimulR (Diaz-Uriarte, 2017), a forward-time genetic simulator specifically developed to represent tumoral evolution. OncoSimulR considers a growing and unstructured asexual haploid population in which mutations can have different effects (none, negative or positive) on cell birth and death rates. Because we are interested in understanding how *dN/dS* correlates with the overall selection pressure on a given genomic region (hereafter, a “gene”), in our simulations, we consider that each cell carries a genome uniquely composed of three genes in which any mutation has none (“neutral gene”), negative (“deleterious gene”) or positive (“driver gene”) effects on fitness, respectively. Note that for our purpose, what matters is that we simulate genes with distinct net implications on tumor growth and that all mutations in the genome interact with each other to determine the overall cell fitness (i.e., they are completely linked as there is no recombination). The order of the different types of mutation along the genome, or even within a gene, or the number of genes considered is not relevant for interpreting these simulations. For simplicity, the only component of fitness we vary is the cell birth rate, i.e., we assume the same death rate for all cells. Each of the three genes encompasses 10,000 biallelic sites, half non-synonymous (*N*) and half synonymous (*S*). In the driver gene, we assume that 0.2% of the *N* sites are functional and have a positive effect on cell fitness (i.e., there is a maximum of 10 driver mutations increasing fitness). In the deleterious gene, we assume that 50% of the *N* sites are functional and have a negative effect on cell fitness. Mutations occurring in the remaining *N* sites and all the *S* sites do not affect cell fitness.

We assume that tumors grow in a continuous-time fashion, following an exponential model defined by the following birth (*b*) and death (*d*) rates:

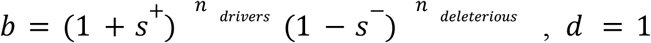

where *s*^+^ and *s*^−^ are, respectively, the selection coefficient of driver (positive) and deleterious (negative) mutations, *n_drivers_* and *n_deleterious_* are, respectively, the number of driver and deleterious mutations in the genome. Each simulation starts with a population of 20 normal cells that divide with *b* = *d* = 1.0, i.e., the population size is initially constant. In each cell division, mutations occur with equal probability at any site that has not previously mutated (i.e., we exclude recurrent mutations) at a rate of *μ* = 10^-6^ mutations per site per cell division. As explained above, depending on the site where the mutation occurs, it will be synonymous or non-synonymous. Then, depending on the gene encompassing this site, it will be a neutral, deleterious, or driver mutation. The selection coefficient for these mutations will vary across scenarios but will be constant in a given tumor for a given type. After the first driver mutation appears, the tumor grows until it reaches an arbitrary threshold of 10^6^ cells (we stop at this size to maintain reasonable computation times). At this point, the simulation stops, and we count the different types of mutations accumulated in the tumor. Note that when a tumor grows very fast, the final population size at which mutations are counted can be slightly larger than this threshold.

### Calculation of dN/dS ratios

To obtain cohort-wide *dN/dS* ratios for neutral, deleterious, and driver genes, we count the number of mutations present at *N* and *S* sites for each gene type across multiple tumors. Given that the number of *N* and *S* sites is the same for the three genes, *dN/dS* is simply the ratio of the number of mutations at *N* and *S* sites. For this calculation, we consider different variant allele frequency (VAF) detection thresholds, for example including only mutations with VAF > 0.01 or > 0.05, or only clonal (fixed) mutations (VAF > 0.95).

### Simulation experiments

We carried two simulation experiments. In the first experiment, where we aim to understand whether the *dN/dS* statistic can correctly identify neutral, deleterious and driver genes, we simulated 100 cohorts of 100 tumors each, resulting from the orthogonal combination of five selection coefficients for driver (*s^+^* = 0.1, 0.25, 0.5, 0.75, 1) and six for deleterious (*s^−^* = −1, −0.75, −0.5, −0.25, −0.1, 0) mutations, using a mutation rate of 10^-6^. In the second experiment, where we assess the relationship between the selection coefficient and *dN/dS*, we carried out a set of simulations with only one gene where all mutations were either deleterious (in 50% of the *N* sites) or driver mutations (in 0.2% of the *N* sites). Here, to describe how this relationship changes throughout time, we also measured *dN/dS* at different time intervals defined by the tumor size (10^3^, 10^4^, 10^5^, and 10^6^ cells).

Under some negative selection scenarios, all mutations may be deleterious, and tumors cannot grow. In these cases, we started the simulation with a population of 1,000 cells and allowed cell division for 50,000 time units. OncoSimulR is a continuous-time simulator, and time units are arbitrary but, under the parameters of our model, a time unit corresponds with the average time for a cell division under neutrality (*b* = 1).

All the scripts to run and analyze the simulations are available at https://github.com/anpefi/pNpS_sims.

## Results

### Average cohort dN/dS for neutral, deleterious, and driver genes

In general, *dN/dS* in a tumor cohort for the neutral gene (Figure 1A, D, G) was, as expected, around 1.0, independently of the strength of selection at the linked deleterious and driver genes and of the VAF threshold. For the deleterious gene, *dN/dS* was invariably 0.5, independently of the strength of selection at the linked driver gene (Figure 1B, E, H). Note that 0.5 is an arbitrary limit imposed by simulating N sites in which only half are functional. If we change this setting to 2%, then *dN/dS* stabilizes at 0.98 (data not shown). At the driver gene, *dN/dS* depended mainly on the effect of the driver mutations (Figure 1C, F, I), with little influence of the linked deleterious mutations. When subclonal mutations are considered in a driver gene, *dN/dS* increases with the strength of selection more or less linearly. However, when *dN/dS* is computed only with the clonal mutations, its magnitude slightly decreases and tends to stabilize in some cases at higher selection coefficients.

**Figure 1.**
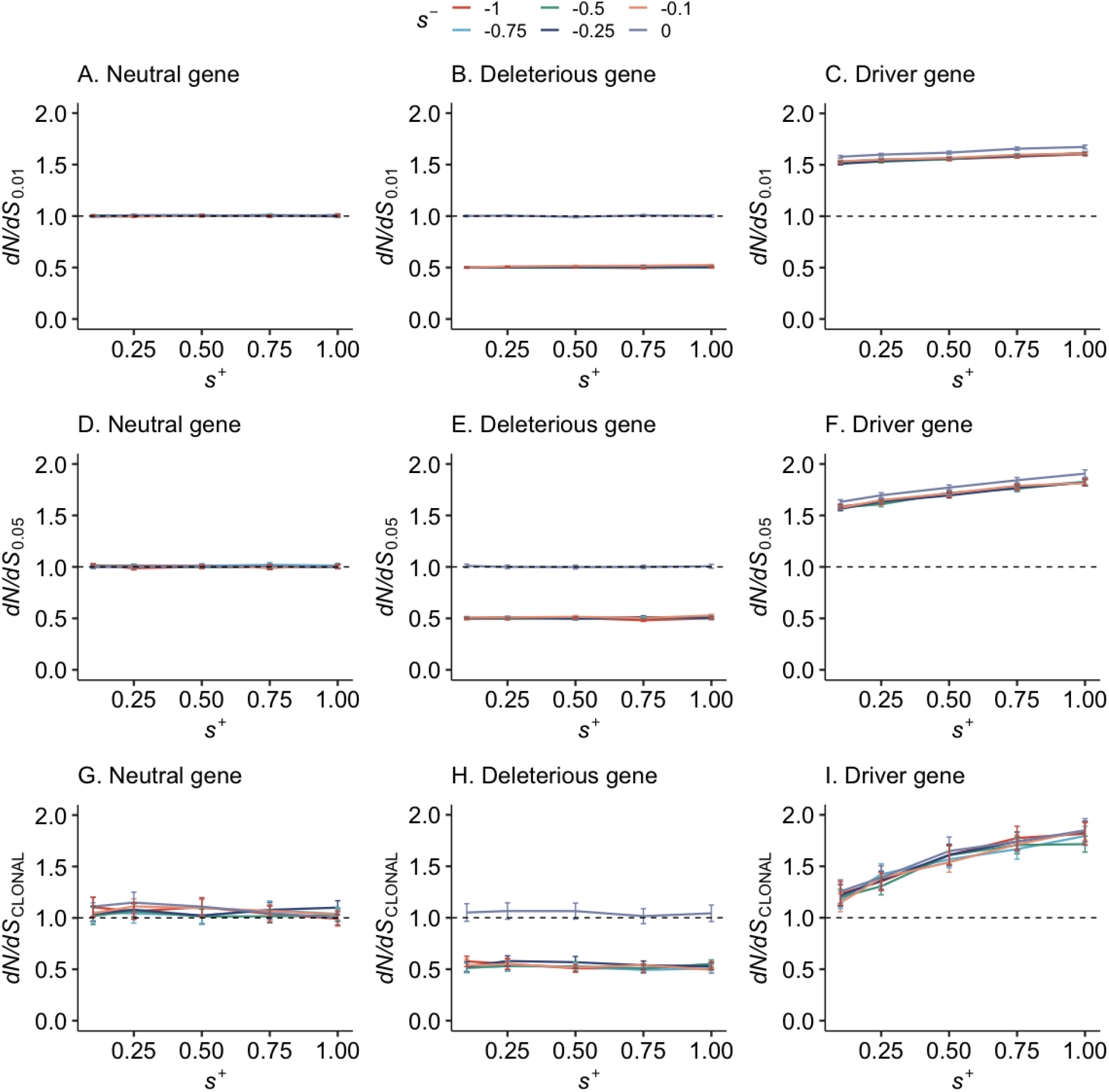
Average cohort-wide *dN/dS* for neutral, deleterious, and driver genes. Positive selection coefficients (s^+^) are represented along the x-axis, while negative selection coefficients (s^-^) are indicated as colored lines. Error bars indicate twice the standard error of the mean.

### Variant allele frequencies at linked neutral, deleterious, and driver genes

To better illustrate the differences between the types and effects of mutations and the allele frequency thresholds, we plotted the reverse cumulative VAF distributions for the nonsynonymous and synonymous mutations for the neutral, deleterious, and driver genes in all the scenarios (Figure 2). In this figure, the trends for a given same mutation type (N or S) only change depending on the strength of selection at the driver gene, so we represent all *s* ≤ 0 by a single line N or S line. The number of mutations decays exponentially with the VAF; however, there is an inflection point around VAF = 0.1 where this decay is reduced. This slowdown of the VAF distribution is caused by selection, and it is more evident with higher selective pressures, particularly in the driver gene.

**Figure 2.**
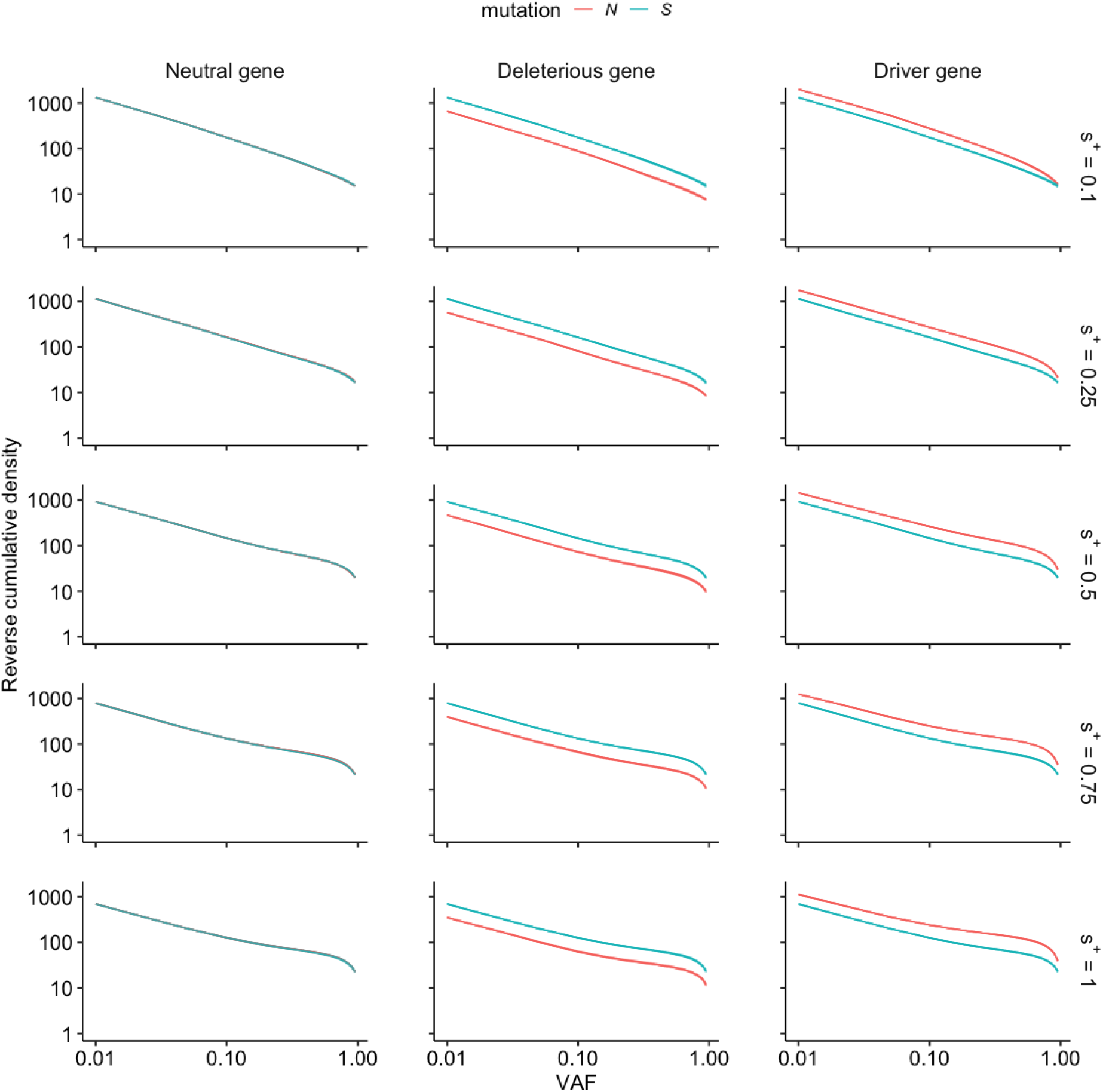
Average distribution of non-synonymous (*N*, red line) mutations and synonymous (*S*, blue line) mutations in blue in all the 100,000 simulated tumors per scenario. The distribution shown is the reverse cumulative density of the number of mutations at a given VAF (both axes in logarithmic scale). Rows correspond to different positive selection and columns to the type of gene. All lines for s ≤ 0 are identical for the same mutation type and therefore are represented by a single red (N) or blue (S) line.

### Relationship between dN/dS and the selection coefficient

To explore in more detail the relationship between *dN/dS* and the selection coefficient *s*, we ran simulations with a single gene in which all mutations had the same selective effect (neutral, negative, or positive). When *s* = 0, *dN/dS* was centered around 1.0, regardless of the detection threshold (Figure 3). When *s* <0, *dN/dS* was more or less constant around 0.5, independently of the VAF threshold. This value derives from the fact that just 50% of the N sites can harbor deleterious mutations. Such mutations are being effectively removed before they reach the detection frequency threshold. On the other hand, when *s* >0, *dN/dS* increased with a more significant selection coefficient and faster for higher VAF thresholds. *dN/dS* values when the frequency threshold was 0.05 or 0.95 (clonal) were very similar.

**Figure 3.**
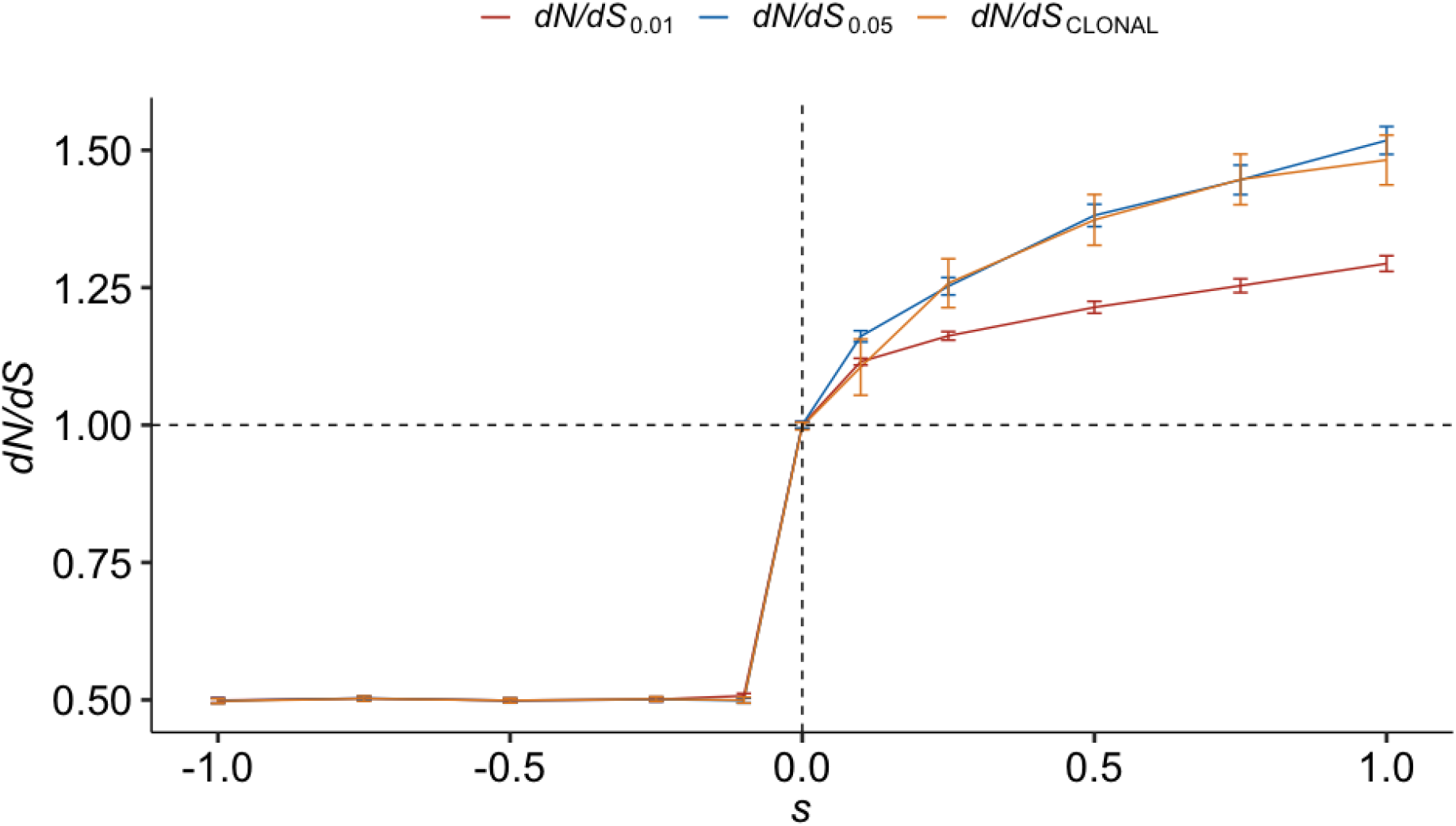
Relationship between the selection coefficient (*s*) and *dN/dS* at different VAF thresholds: VAF > 0.01 (red), VAF > 0.05 (blue), VAF > 0.95 (clonal mutations, orange). Lines show the mean value across cohorts, and error bars indicate twice the standard error of the mean.

To better understand these trends, we represented the reverse cumulative VAF distributions of *N* and *S* mutations for the single-gene scenarios along with tumor progression for different positive selection scenarios (Figure 4). Here we can see an inflection point between 0.05 and 0.1 VAF that becomes more abrupt as the tumor grows. Given enough time (>10^4^ cells), there is a stabilization due to the action of selection.

**Figure 4.**
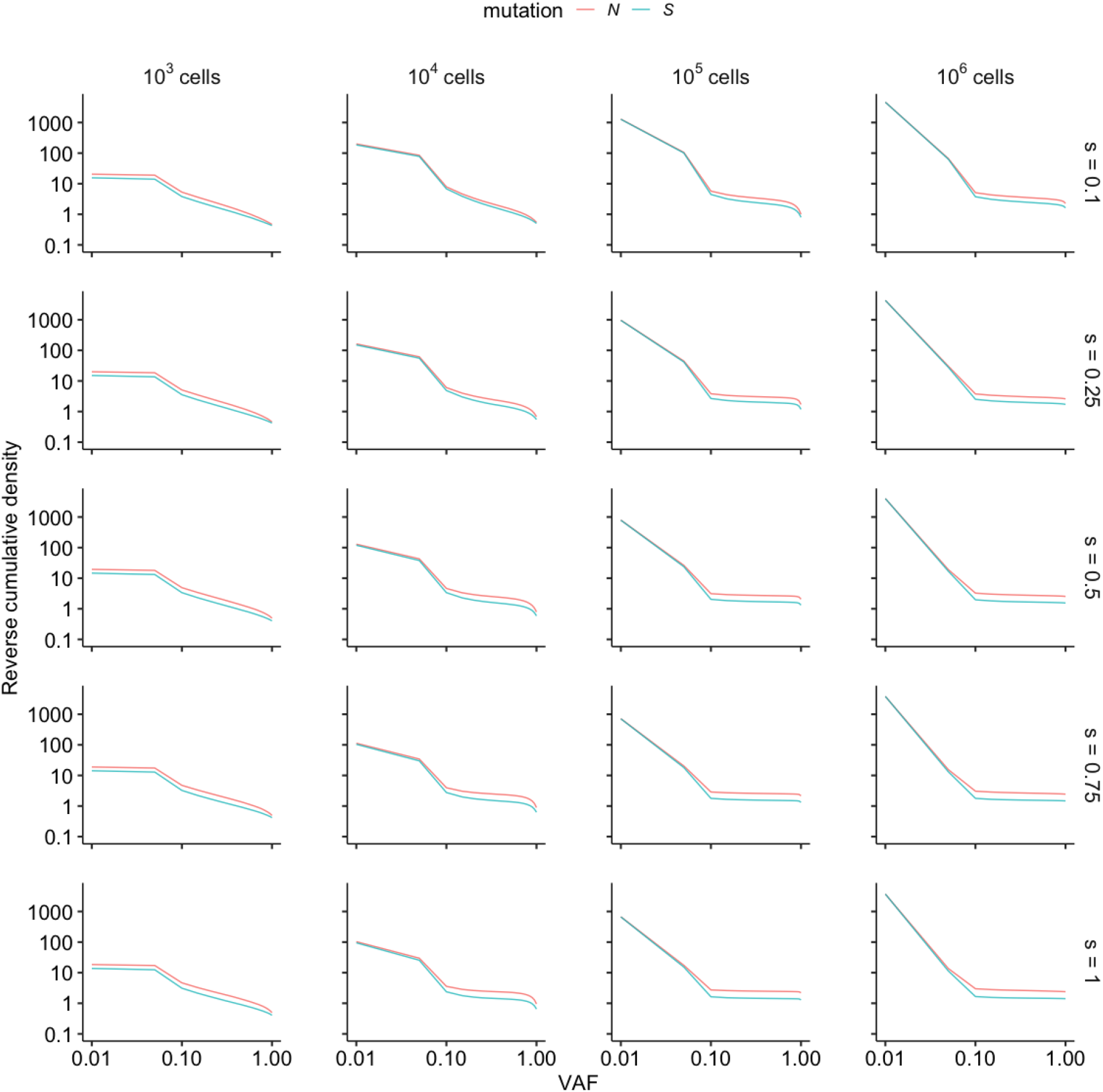
Average distribution of non-synonymous (*N*, in red) mutations and synonymous (*S*, in blue) mutations in blue in all the 100000 simulated tumors per scenario. The distribution shown is the reverse cumulative density of the number of mutations at a given VAF (both axes in logarithmic scale). Rows correspond to different positive selection and columns to different sampling times.

### Temporal dynamics of dN/dS in a growing tumor

To study the relationship between *dN/dS* and the selection coefficient along with tumor growth, we measured *dN/dS* at different tumor sizes. In general, *dN/dS* grows over time (Figure 5). However, if we include subclonal mutations in the calculation of *dN/dS*, there is a slight mismatch in the early stages where the initial appearance of driver mutations eventually raises the *dN/dS* ratio, which rapidly decreases as the first driver mutations are fixed (Figure 5A, B). Once the initial driver mutation is fixed, additional driver mutations will inflate the *dN/dS*, particularly at higher selection coefficients. When computing *dN/dS* only with the clonal mutations, with high selection pressure, *dN/dS* seems to stabilize after the tumor reaches 10,000 cells (purple line in Figure 5C). This plateau is more apparent when selection is strong and tumor growth is fast. Then, only one, on average, driver mutation has become clonal before the simulation finishes.

**Figure 5.**
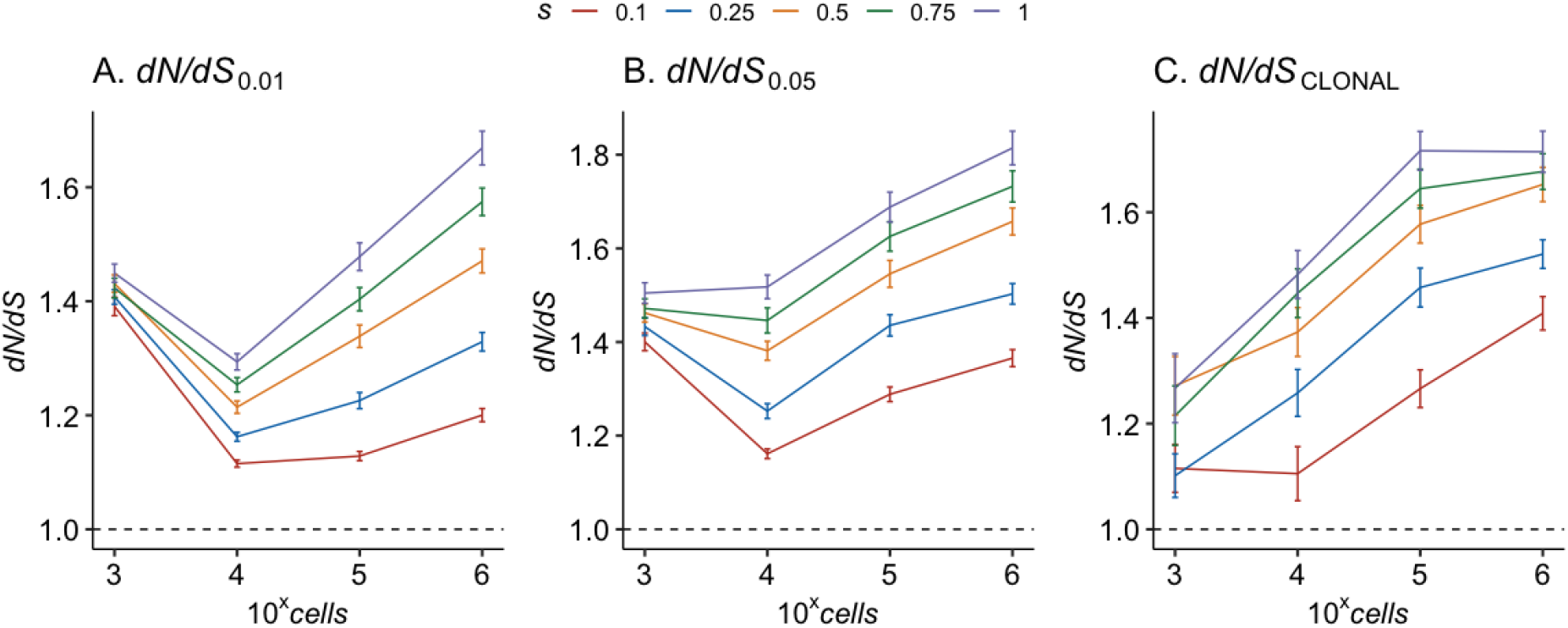
Evolution of the *dN/dS* ratio along tumor progression under different positive selection coefficients (*s* from 0.1 to 1, color lines) using three different VAF thresholds (0.01, 0.05 and 0.95 (CLONAL)). Lines show the mean value for all the cohorts, and error bars indicate twice the standard error of the mean.

## Discussion

The ratio between the rate of non-synonymous substitutions per non-synonymous site and the rate of synonymous substitutions per synonymous site, *dN/dS* has been actively used to identify evolutionary pressures in cancer. However, *dN/dS* was originally developed for diverged genomes, where differences represent fixed mutations rather than segregating polymorphism, as is the case for tumors. Moreover, for samples taken from asexual populations, like cancer cells, the relationship of *dN/dS* with the selection coefficient can be distinct to the one assumed for intraspecific comparisons, biasing the inference of selective pressures from *dN/dS* (Kryazhimskiy and Plotkin, 2008). In asexual cell populations, where all sites are effectively linked, the value of *dN/dS* changes with the allele frequency detection threshold (Williams et al., 2020). Therefore, it is interesting to understand to which extent *dN/dS* can be used to reliably identify driver, neutral or deleterious events in cancer genomes and its relationship with the selection coefficient under different evolutionary regimes.

Using computer simulations, here we show that we can effectively use *dN/dS* to distinguish whether a genomic region (i.e., a “gene”) mutated in a cancer cohort has an overall neutral, driver, or deleterious role, even when it is linked to genes under opposite selective pressures. On the other hand, *dN/dS* is not sensitive to the strength of selection under some scenarios. Around 97-98% of the genes along cancer genomes show *dN/dS* values around one and, therefore, are deemed neutral (Martincorena et al., 2017). In our simulations, neutral genes, despite linkage with genes under negative and positive, result in *dN/dS* values always around one.

There is some controversy around negative selection acting on cancer. Martincorena et al. (2017) suggest that negative selection on point mutations is absent during tumor progression and healthy somatic tissue maintenance. Tilk et al. (2019) argued that the absence of negative selection in cancer is due to the poor efficacy of selection in non-recombining regions (i.e., a Hill-Robertson effect). On the other hand, Zapata et al. (2020, 2018) infer negative selection on essential and activating immune response genes. These studies have in common that they used the *dN/dS* to detect selection. While this debate continues, our simulations show that *dN/dS* can be used in tumor cohorts to identify genomic regions under overall negative pressure, even if they are linked with other genomic regions under strong positive selection, as in cancer.

Genes with an overall *dN/dS* above 1 in tumor cohorts are considered driver genes. Our simulations suggest that, despite some of the caveats about using this metric within populations (Kryazhimskiy and Plotkin, 2008), this interpretation is typically correct, even in the case of cancer, where genes under divergent selective pressures might coexist in the same genome. However, if only clonal mutations are used for the computation, *dN/dS* might not be sensitive to significant positive selection coefficients. This is because, except for the earliest ones, additional driver mutations find it more challenging to fixate and stay at subclonal frequencies.

The point in time in which we examine a polymorphism for signs of past selective pressures matters. Our results suggest that hitchhiking can occur very early in tumor evolution, producing a slight excess of mutations at high frequencies with respect to what is expected only by genetic drift. This leads to *dN/dS* growing over time since the new mutations that escape the effect of drift, and reach the VAF threshold, are being swept by those with a positive effect. In other conditions, without growth and with free recombination, *dN/dS* shows a phase of change (up and down) until reaching stationarity, depending on the time of divergence of the sequences and therefore on the relative amount of polymorphisms and substitutions (Kryazhimskiy and Plotkin, 2008; Mugal et al., 2020, 2014). In our model, clonal mutations are more likely to be represented by early driver, and passenger mutations hitchhiked along. Therefore, when *dN/dS* is computed only with clonal mutations and selection is strong, it tends towards stabilization over time, as few other mutations reach clonality. In this case, dN/dS reflects the initial selection pressure on the tumor.

On the other hand, if we lower the frequency threshold to include subclonal mutations, *dN/dS* increases over time because more recent driver mutations at intermediate frequencies are then considered. Overall, *dN/dS* will depend on the amount of polymorphism segregating at sampling time. This agrees with the need to correct for mutation frequency when estimating fitness effects in somatic evolution (Williams et al., 2020).

Finally, we cannot forget that given that tumor biopsies typically include a tiny proportion of the tumor cells and the pervasive levels of intratumoral heterogeneity observed, most empirical datasets are subject to sampling bias (Chkhaidze et al., 2019). This means that apparently, clonal mutations might still be segregating in the population.

### Conclusions

We have used a simple model of cancer evolution under different selective pressures to elucidate the combined effects of negative and positive selection on the *dN/dS* ratio typically measured in cancer genomics. We have shown that, even if *dN/dS* is designed for divergent populations that have accumulated fixed differences, we can use it in cancer cohorts to identify genes under overall negative and positive selection.

## Acknowledgments

This work was supported by project No. 32030, co-financed by COMPETE 2020, Portugal 2020, and the EU through the ERDF, and by FCT (PTDC/BIA-EVL/32030/2017, assists A.P.F) through national funds, by the European Research Council (ERC-617457-PHYLOCANCER awarded to D.P.) and by the Spanish Ministry of Science, Innovation, and Universities - MCIU (PID2019-106247GB-I00 awarded to D.P.). D.P. receives further support from Xunta de Galicia.

